# EPIGENETIC LANDSCAPE OF HEAT STRESS INTERGENERATIONAL INHERITANCE IN A TELEOST FISH

**DOI:** 10.1101/2022.10.17.512480

**Authors:** Aurélien Brionne, Anne-Sophie Goupil, Stéphanie Kica, Jean-Jacques Lareyre, Catherine Labbé, Audrey Laurent

**Affiliations:** INRAE, UR1037 LPGP, Fish Physiology and Genomics, Campus de Beaulieu, F-35000 Rennes, France

**Keywords:** Spermatozoa, DNA Methylation, Temperature, Inheritance, Fish

## Abstract

Epigenetic information is transmitted from cell to cell, and even generation to generation. The question of epigenetic inheritance in fish has become of crucial interest in the recent years, when the mammalian model of methylome erasure in germ cells and embryos was found not to be conserved. Fish, which are particularly exposed to environmental variations might thus be prone to transmit epigenetic alterations to their offspring, driving rapid environmental acclimation. Here, by sequencing spermatozoa and muscle methylomes, we characterized the methylation landscape of paternal gametes in rainbow trout and demonstrated its sensitivity to a 4°C increased rearing temperature during spermatogenesis. We found that spermatozoa methylome primes housekeeping and developmental genes for activation and might be instrumental to early development. Most of these methylation-free promoters were not affected by temperature, attesting the robustness of the epigenetic programming of early development. However, the increase of temperature triggered the differential methylation of 5,359 regions, among which 560 gene promoters control spermiogenesis and lipid metabolism. We therefore report, for the first time in fish, that sperm epigenetic landscape carries marks of parental environmental conditions. In the context of a 4°C temperature increase during spermatogenesis, we describe how rainbow trout sperm DNA methylation might be a molecular basis of intergenerational inheritance and question its role in controlling next generation’s performances and acclimation to climate change.

## INTRODUCTION

Epigenetic landscapes define cell identities by regulating chromatin 3D architecture and transcriptional programs. They rely on DNA methylation and histone post-translational modifications, and are either inherited from a mother cell or modulated upon physiological context. DNA methylation is the addition of methyl group to the 5^th^ or 4^th^ position of a cytosine: 5-methylcytosine (5mC) or *N*^4^-methylcytosine (4mC), respectively, or the 6^th^ position of an adenine: *N*^6^-methyladenine (6mA) (Anton and Roberts, 2021). DNA methylation appeared in prokaryotes, in a restriction-modification system believed to act as a primitive immune defense mechanism. Cells that possessed DNA MethylTransferase enzymes were able to distinguish self from non-self DNA when infected by non-methylated exogenous DNA (Suzuki and Bird, 2008). DNA methylation spread through evolution and is found in almost every species from fungi and plants to mammals. The most common form of genomic DNA methylation in animal Eukaryotes is 5mC. In contrast to bacteria and archaea in which it can be found in various genetic contexts such as CpHpG or CpHpH (H referring to either A, C or T), 5mC is mostly detected in the symmetrical CpG dinucleotide context in vertebrates. Of all kinds of DNA methylation, 5mCpG is therefore the most abundant and most studied form in in these species. Interestingly, genomic 5mCpG profiles evolved in a clade-specific manner (Aliaga et al., 2019), possibly reflecting different use of methylation during evolution. While most vertebrates exhibit global DNA methylation, spanning all over the genome at the exception of regulatory elements, some model species such as *C. elegans* or *D. melanogaster* possess virtually no 5mC. Alternatively, other species such as *Crassostrea gigas* or *Api mellifera* exhibit mosaic-type of methylation, with large genomic regions being highly methylated interspersed with methylation-free regions. A conserved function of DNA methylation in most eukaryotes is repeats and transposable elements silencing, which might be inherited from its primary role in parasitic DNA control. In addition, it contributes to transcriptional regulation by modulating the binding of transcription factors to their target sequences, or by interacting with methyl-CpG-binding proteins in turn able to recruit, in turn, chromatin remodelers (Greenberg and Bourc’his, 2019). Of note, 5mC was not only maintained through evolution, it also contributed to genomes expansion and innovation, by allowing the integration and assimilation of transposable elements (Zhou et al., 2020).

As part of an interface between genome and environment, the DNA methylation is susceptible of modulation by methylation/demethylation machineries upon various endogenous or exogeneous stimuli, such as development and aging, cancer, viral infection, or environmental change. A famous example of water temperature effect on fish DNA methylome is the modulation of the aromatase promoter leading to a sex reversal from genetic female to phenotypic male individuals (Navarro-Martín et al., 2011). In vertebrates, 5mC deposition is mediated by DNA MethylTransferases (DNMT) which catalyze the addition of a methyl group from the S-adenosylmethionine (SAM) donor metabolite to cytosines and are of two types. Following replication, maintenance DNMTs are recruited on hemi-methylated DNA and copy the methylation pattern on the newly synthetized DNA strand. On the other hand, *de novo* DNMTs are able to bind and methylate so far non-methylated DNA regions. The erasure of DNA methylation marks can occur in a cell as the result of proliferation under inhibition of a maintenance DNMT (passive demethylation), or active demethylation *via* the iterative oxidation of 5mC by Ten Eleven Translocation (TET) enzymes and consecutive activation of the Thymine DNA Glycolsylase (TDG)-dependent base excision repair (BER) pathway. DNA methylation patterns are therefore concomitantly inheritable through cell division and reversible during the life of a cell.

Importantly, increasing evidence indicate that epigenetic information, and *a fortiori* its alterations, are also transmittable through meiosis and fertilization in fish, from generation to generation. This is particularly suggested by environmental acclimation studies in fish, where offspring performances were shown to be influenced by the life conditions of their genitors. In the tropical reef fish *Acanthochromis polyacanthus*, the aerobic scope of F2 individuals raised from hatching to adulthood at higher temperatures (+1.5°C and 3°C) is increased if their F1 parents were also reared at these increased temperatures (Donelson et al., 2012). In the stickleback *Gasterosteus aculeatus*, eggs hatching success and embryo size are influenced by both maternal and paternal exposure to increased temperature (Shama and Wegner, 2014). In the tongue sole *Cynoglossus semilaevis*, thermally-induced sex reversed F1 pseudo-males have sex-reversed pseudo-male offspring although these F2 larvae are not exposed to thermal induction (Chen et al., 2014). In the rainbow trout *Oncorhynchus mykiss*, offspring thermotolerance (survival and growth) is improved when genitors also have experienced increased rearing temperature during gametogenesis (Butzge et al., 2021). These data argue in favor of an intergenerational epigenetic inheritance, and suggest that gametes can transmit environmental acclimation clues to the offspring. Very few studies investigated fish gametes epigenetics. However, Fellous *et al*. reported the differential expression of various epigenetic modifiers in gonads and gametes upon increased temperature in stickleback, suggesting that gametes might be responsive to temperature variations (Fellous et al., 2022). In rainbow trout, Gavery *et al*. found differentially methylated DNA regions in spermatozoa from males of wild versus hatchery origins (Gavery et al., 2018), further indicating that paternal gametes epigenome might be modulated by the environment. Literature thus suggests that sperm DNA methylome can be sensitive to environmental conditions and therefore might be a molecular vector of paternal intergenerational epigenetic transmission in fish. In fact, the importance of zebrafish sperm DNA methylome for offspring epigenetic programming was demonstrated by Potok *et al*. (Potok et al., 2013) and Jiang *et al*. (Jiang et al., 2013) who revealed that embryo DNA methylome is inherited from spermatozoa, maternal DNA methylome being remodeled before zygotic genome activation to match the paternal one. The role of sperm DNA methylome for zebrafish embryo development was further emphasized by the demonstration that it drives the gradual increase of chromatin accessibility and gene activation during zygotic genome activation (Liu et al., 2018). Ortega *et al*. demonstrated that zebrafish, in contrary to mammalian models, do not undergo global DNA methylation reprogramming of primordial germ cells, unraveling the persistence of the epigenetic memory through generations in the species and a potentially important role of gamete DNA methylation in heredity and evolution (Ortega-Recalde et al., 2019). Unfortunately, fish gametes epigenomic studies are scarce and mostly limited to zebrafish. For a better understanding of the molecular basis underlying fish intergenerational inheritance, data reflecting the full diversity of fish species and reproduction modes are missing, and particularly in the omics perspective.

Given the potential importance of the paternal methylome in fish and in the context of environmental change, we sought to characterize the rainbow trout *(O. mykiss)* sperm methylome and its sensitivity to a water temperature increase. To this aim, we performed whole genome bisulfite sequencing (WGBS) on sperm DNA of trout reared at 12°C or 16°C during gametogenesis. Muscle samples from fish raised at 12°C were sequenced as somatic reference tissue. We used these single-base resolution data sets to (i) characterize genome-wide methylation dynamics in representative rainbow trout germline and somatic tissues, (ii) highlight the paternal gamete methylome specificities, (iii) assess its sensitivity to a water temperature increase.

## RESULTS

### Rainbow trout genome features and single-base resolution DNA methylomes of sperm and muscle

We briefly summarize here the main features of publicly available rainbow trout genome, which are susceptible to impact DNA methylation. *Oncorhynchus mykiss* genome is made of 2.2 Gb parsed into 29 chromosomes with 41,365 and 5,977 coding and non-coding genes, respectively (according to ensembl v105.1 genome annotation of Omyk_1.0 version by Full genebuild). The guanine and cytosine (GC) content of 43.4 % is relatively high in comparison to 36.6 % found in zebrafish, 40.8 % in medaka, 40.2 % in xenopus, 40.4 % in human or 42.7 % in mouse according to NCBI genome database. The rainbow trout genome contains 70,672,452 CpGs (70,671,452 nuclear; 1,118 mitochondrial), with a global genome-wide observed over expected (o/e) CpG ratio of 0.8, indicative of a slight CpG erosion during evolution.

In this work, we performed whole genome bisulfite sequencing (WGBS) of muscle and sperm (two experimental conditions) DNA samples. WGBS yielded a minimum of 300 million paired-end reads of 150bp per sample (**Tab 1**). After trimming, only uniquely mapped reads on the *O. mykiss* genome or its *in silico* bisulfite converted version were kept for further analysis (using Bismark bio-informatic software and Omyk_1.0 genome version, (Felix Krueger and Andrews, 2011)). In all sample types, the average mapping efficiency of the trimmed reads was of approximately 60%. Our final data set contains 65,125,263 CpGs, representing 92% of the trout genome, 70 to 79 % of them being covered at least 5 x. On average, we obtained a sequencing depth of 10-11 × per strand.

**Tab 1.**
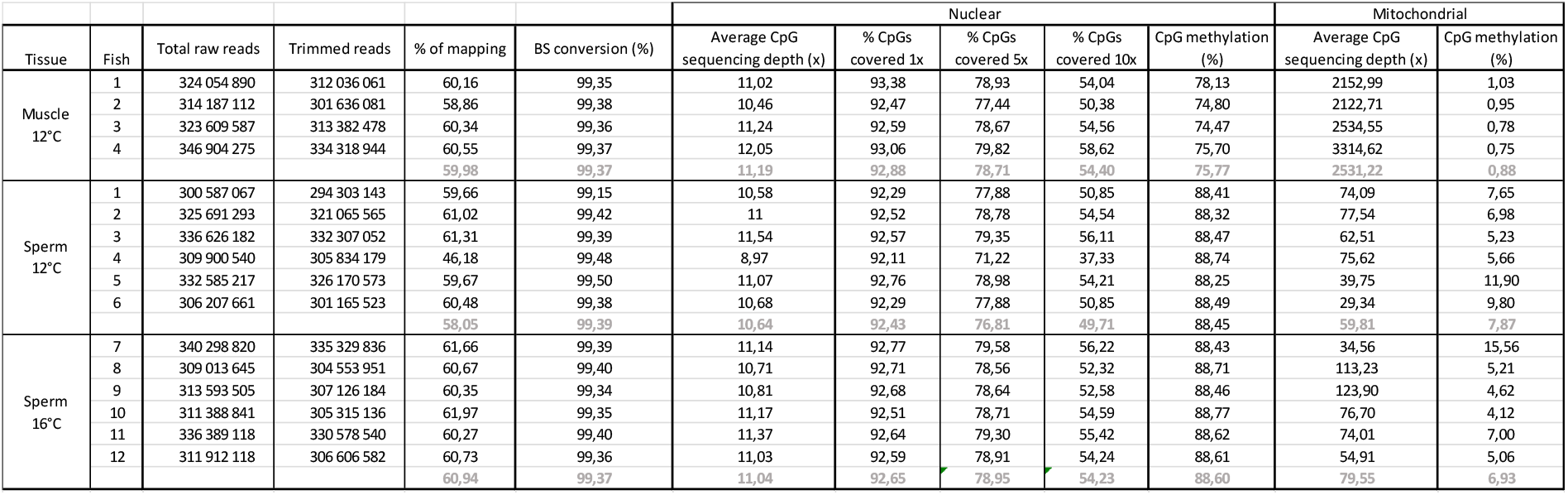
Summary of WGBS. Paired reads were mapped uniquely to the rainbow trout genome (Omik_1.0) using Bismark. Average sequencing depth indicates the mean number of reads per position in nuclear and mitochondrial genomes. The nuclear coverage represents the proportion of mapped CpGs over the total number of CpG sites in the nuclear genome, here calculated under 1X, 5X or 10X sequencing depth selection threshold.

We found that almost all methylated cytosines sit in CpG dinucleotides. Indeed, while we measured average methylation of respectively 74.50 and 87.10 % in CpG context in muscle and sperm, respectively (**Tab 1**), methylated CHG and CHH (H representing A, T or C) were detected at rates lower than 1% (**Suppl Tab 1**). The odd value found for sperm of fish 4 might be due to a technical bias on this sample. Alternatively, it might reflect a poor reliability of CHG and CHH methylation counts related to a low coverage effect on rare events. For this reason, this study is dedicated to the analysis of CpG methylation only. Finally, we detected very low CpG methylation on mitochondrial DNA, both in muscle and in spermatozoa (**Tab 1**). Our sequencing depth analysis revealed an important mitochondrial DNA over-representation in muscle *versus* sperm (**Tab 1**). This agrees with the expected differences in ratios of nuclear to mitochondrial DNAs between these tissues with abundant mitochondria in muscle while trout spermatozoa possess a unique semicircular mitochondrion.

Finally, in order to choose a strategy for the bio-informatic analysis, we wondered whether we should filter out the CpGs suffering low coverage. We therefore tested for a potential bias originating from sequencing depth differences between CpGs. **Supplementary Table 2** recapitulates, for each sample, the number of cytosines in CpG context, being sequenced as methylated in at least one read. When considering the whole data sets, *circa* 93 % of CpGs, in both tissues, showed at least one positive methylation call. Remarkably, these numbers decreased as the coverage threshold applied to the data increased (5 ×; 10 ×), and they dropped faster in muscle than in spermatozoa. In fact, among the cytosines covered at least 10 times, 67 % of them, on average, were sequenced as methylated in at least one read in muscle, *versus* 85 % in spermatozoa. This very reproducible observation among fish samples reveals a fragment selection bias in the sequencing library preparation process, likely reflecting differences in the chromatin organization between tissues. Non-methylated DNA fragments are more likely to lie in open and accessible chromatin, more prone to be resuspended in solution and to be represented in the sequencing libraries. Both muscle and spermatozoa WGBS libraries show a better representation of non- or poorly-methylated CpGs, but interestingly, this bias is much milder in spermatozoa, reminiscent of a less heterogeneous chromatin conformation in this cell type. We therefore concluded that lowly covered CpGs are informative and decided not to apply any filter on sequencing depth. Data were instead analyzed using DSS bio-informatic tool and its optional smoothing of methylation levels on 500 bp windows (Feng et al., 2014; Wu et al., 2015). This allows the compensation of poorly sequenced CpGs by the surrounding ones.

### Rainbow trout genome is globally methylated both in muscle and spermatozoa

To our knowledge, we present the first comprehensive rainbow trout methylomes of somatic and germline representative cell types. We therefore questioned the general trout DNA methylation pattern in a descriptive analysis of muscle and sperm coupled WGBS data sets (**Suppl Fig 1, fish 1 to 4**). We first assessed the methylation profile on various genomic features, by calculating the average methylation rates along a reconstituted metagene (**Fig 1A**), around transcription start sites (TSS) and transcription end sites (TES) (**Fig 1B**) of 115,853 annotated Ensembl coding and non-coding transcripts (ensembl v104.1, Omyk_1.0). As already suggested in **Tab 1**, we observed a global DNA hypermethylation of sperm compared to muscle. This result is consistent with the frequently reported high methylation level of spermatozoa in vertebrates (Jiang et al., 2013; Kobayashi et al., 2012; Potok et al., 2013). Nevertheless, both tissues showed a drastic drop of CpG methylation on proximal promoter regions (1kb upstream of TSS) while gene bodies exhibited particularly high methylation rates. This was further emphasized in the analysis of the distributions of average methylation in various functional elements (**Fig 1C**). We found that the methylation rates of promoter regions, 5’UTR, first exon and first intron segregated in a clear bimodal mode, being either hypomethylated (< 15 % methylation) or heavily methylated (> 60 % in muscle, > 80 % in sperm). On the contrary, nearly 100 % of other exons, introns, 3’UTR and downstream regions were uniformly highly methylated, with higher ratios in spermatozoa than in muscle (**Fig 1C**). It is important here to stress that CpG methylation rates are calculated as a mean of signals coming from individual cells. Therefore, the more homogeneous is a cell type, the more methylation counts are expected to be close to either 0 % or 100 %. In spermatozoa, which are a fully differentiated and non-proliferating, intercellular variability is in fact very low, and methylation rates are more discrete than in muscle. Finally, we sought to interrogate methylation rates in repeat sequences. Repeat elements identified using RepeatMasker (Smit, 2015) were all found to be methylated in both tissues. Transposons, retroposons, LTR, LINEs and SINEs were particularly highly methylated in sperm. Therefore, our analysis of one somatic and one germline tissue showed that the rainbow trout DNA methylation pattern is similar to that of other described vertebrates, exhibiting a typical global methylation in both tissues with the exception of a subset of methylation-poor promoters. Rainbow trout gene bodies also carry high methylation rates already reported in model vertebrates to modulate alternative splicing or prevent spurious transcription initiation. Setd2 enzyme indeed interacts with elongating RNA polymerase II and deposits H3K36me3 on transcribed gene bodies. Dnmt3b is in turn recruited by H3K36me3 and methylates DNA (Neri et al., 2017).

**Fig 1.**
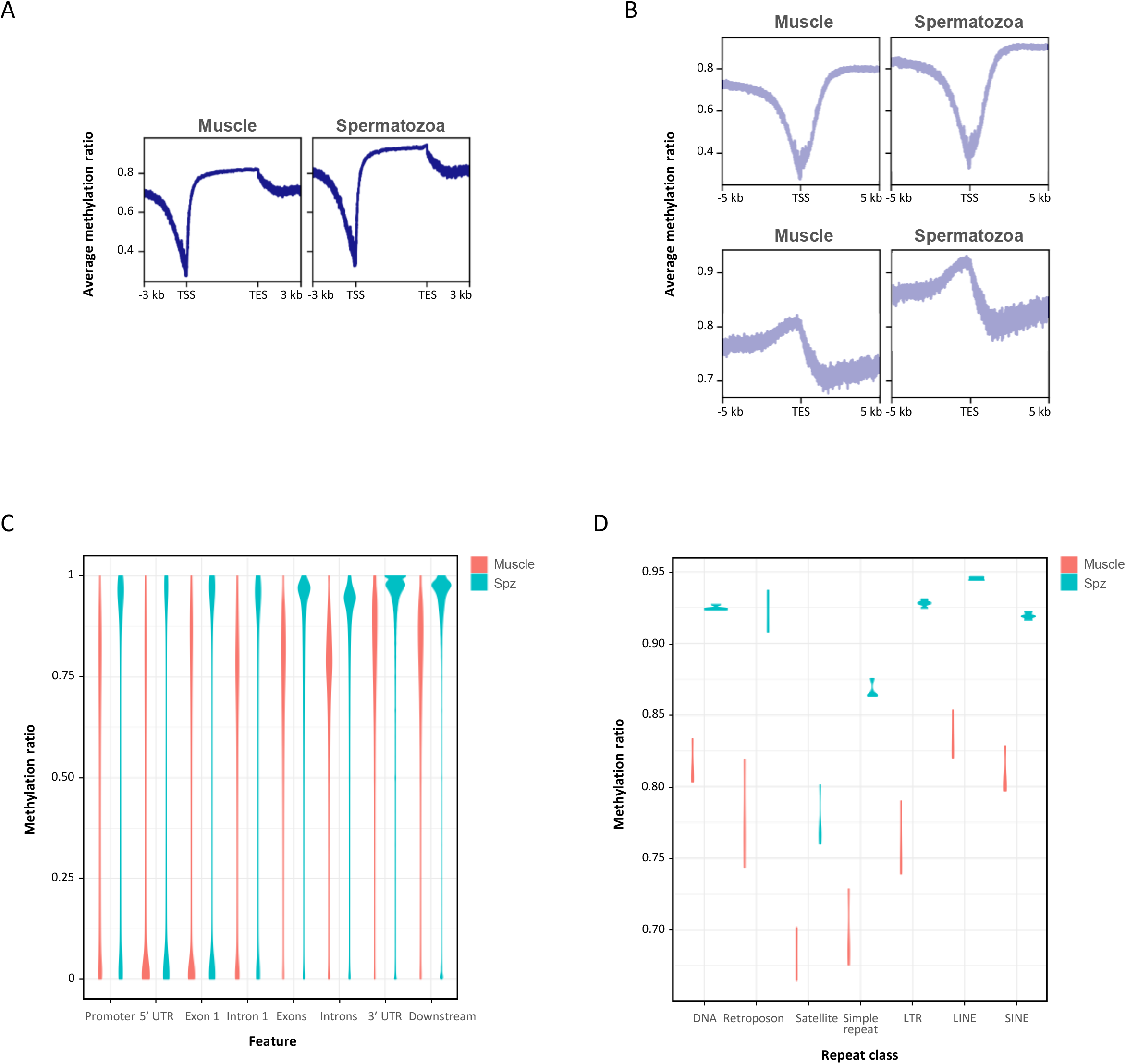
Global DNA methylation in rainbow trout muscle and spermatozoa according to genomic features. A. Average methylation profiles along the 115 853 annotated ensembl coding and non-coding transcripts of ensembl v104 Omyk_1.0 genome. B. Average methylation profiles around TSS (+/− 5 kb) (left panels) and TES (+/− 5 kb) (right panels). C. Distribution of methylation ratios according to gene features: promoter regions (−1 kb; TSS), 5’UTR, exon1, intron1, all exons, all introns, 3’UTR and downstream regions (TES; +1 kb). D. Distribution of methylation ratios found in muscle and sperm in transposons (DNA), retroposons, satellites, simple repeats, LTR containing elements, LINEs and SINEs.

### The signature of sperm methylation profile reflects spermatogenesis and embryonic development

Seeking for spermatozoa methylome specificities we then ran a comparative analysis of sperm and muscle coupled WGBS data (**Suppl Fig 1B, fish 1 to 4**). A first approach, by principal component analysis (PCA), led us to observe a clear segregation of sperm and muscle samples in spite of a greater dispersion of muscle samples, likely reflecting the greater complexity of this multicellular tissue compared to sperm (**Fig 2A and Suppl Fig 2**).

**Fig 2.**
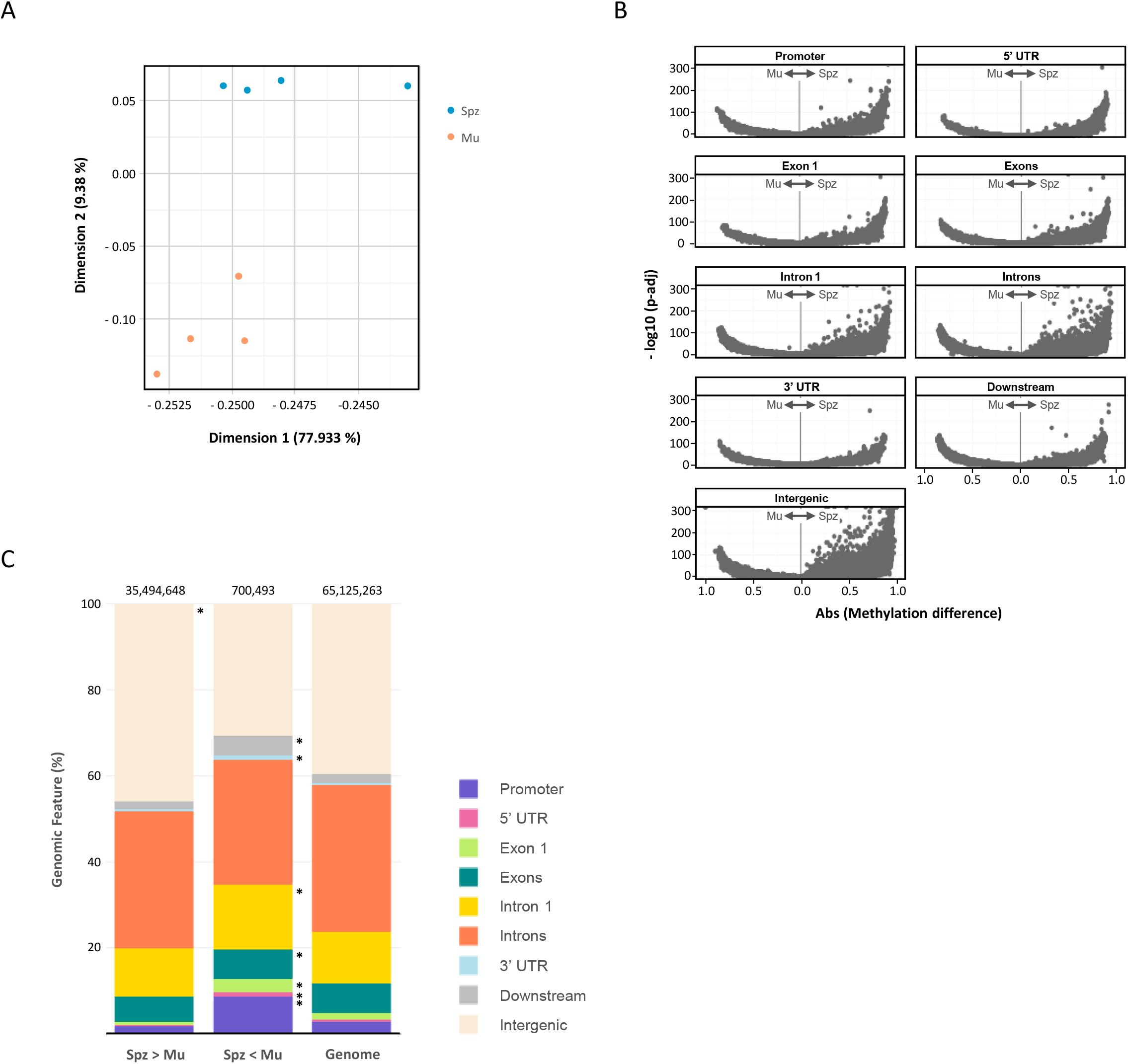
Differentially methylated cytosines between muscle and spermatozoa. A. PCA plot of muscle and spermatozoa WGBS data sets. B. Volcano plots of DMCs between muscle and spermatozoa in every genomic feature (× axis: absolute methylation difference, y: −log10 (adusted p-value)). C. Relative distributions of DMCs (Spz > Mu: hypermethylated in sperm compared to muscle; Spz < Mu: hypomethylated in sperm compared to muscle) according to genome features (promoter regions (−1 kb; TSS), 5’UTR, exon1, intron1, all exons, all introns, 3’UTR, downstream (TES; +1 kb) and intergenic regions). The reference set (Genome) contains all analyzed CpGs. Genome features statistically enriched in DMCs compared to the reference set are indicated with an * (binomial test).

Differentially meythylated CpGs (DMCs) between muscle and spermatozoa were identified by DSS (Feng et al., 2014; Wu et al., 2015). We found 36,195,141 DMCs with a threshold adjusted p-value of 0.05, meaning that approximately half of the genome is differentially methylated between muscle and spermatozoa. These DMCs are spread over every chromosome (not shown) and across all genomic features (**Fig 2B, C**). As expected, most of DMCs (98 %) are hypermethylated in sperm compared to muscle. Interestingly, while DMCs hypermethylated in spermatozoa are preferentially found in intergenic regions (also representing the largest part of the genome) (**Fig 2B**), the few sperm-specific hypomethylated DMCs, are significantly enriched in gene bodies and promoters. This is indicative of a structural role of sperm DNA hypermethylation on whole genome chromatin. On the contrary, the enrichment, in spermatozoa, of hypomethylated DMCs in genes and gene-proximal sequences suggests that a few promoters specifically escape the global methylation and might be instrumental for the proper execution of spermatogenic transcriptional program.

In order to gain insight into this spermatozoa-versus-muscle-specific transcriptional regulation, we isolated promoters and compared their methylation levels in both tissues. Promoter methylation status is indeed informative of a transcriptionally repressed or permissive state (however still sensitive to histone post-translational modifications). We therefore wondered whether we could evidence muscle- or spermatozoa-specific methylome signatures, which would reflect their functional specificities. Beyond this, our aim was to question whether transcriptionally silent trout spermatozoa still carry marks of spermatogenesis and/or might possess already primed embryonic development genes as expected from previous studies in zebrafish (Lindeman et al., 2011). Surprisingly, the density plot of average methylation levels along promoters shows a remarkable correlation between muscle and spermatozoa (**Fig 3A**): among the 115,853 tested promoters, more than 95% are similarly methylated in both tissues. More precisely, 46 % of them were hypermethylated in both muscle and spermatozoa (methylation ≥ 0,75 %), while 30 % were hypomethylated (methylation ≤0,25 %) in both tissues. However, several gene promoters did show tissue-exclusive behaviors. In order to identify differentially methylated promoters and their biological functions, we split genes into different categories according to their muscle-versus-spermatozoa methylation status and looked for GO term enrichment in each class (**Suppl Tab 3A-D**).

**Fig 3.**
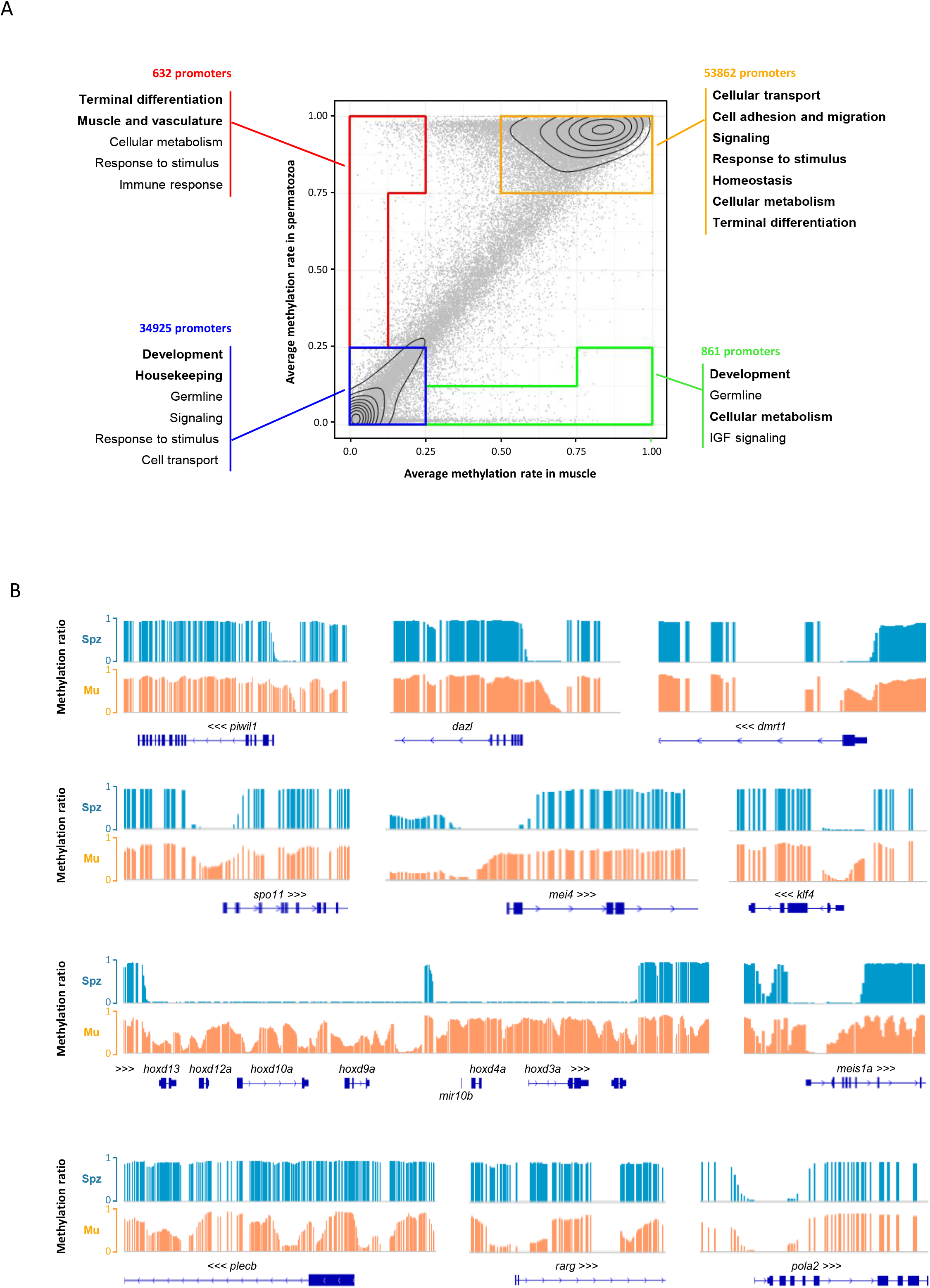
Differentially methylated gene promoters between muscle and spermatozoa. A. Plot of average methylation of ensembl omyk_1.0 v105 transcrits promoters in muscle (x axis) *versus* spermatozoa (y axis). Common hypo-methylated promoters are highlighted in the blue square, common hypermethylated promoters in the orange shape, muscle-specific hypo-methylated promoters in the red shape and spermatozoa-specific promoters in the green shape. Functional annotation terms deduced from a GO term enrichment analysis and corresponding to each class are indicated. B. IGB snapshots of transcripts or TSS regions as examples of spermatozoa-specific hypo-methylated promoters *(piwil1, dazl, dmrt1, spoll, mei4, hox* cluster and *mir10b, meis1a*), muscle-specific hypo-methylated promoters *(plecb* and *rarg*) or common hypomethylated promoter (*pola2*). Muscle: Mu; Spermatozoa: Spz.

Common hyper-methylated promoters (53862 transcripts, **Fig 3A**, orange rectangle, **Suppl Tab 3A**) were particularly enriched in GO terms referring to cellular transport, cell adhesion and migration, signaling, response to stimulus, homeostasis, metabolism and terminal differentiation (angiogenesis, pulmonary valve differentiation, melanocyte differentiation).

On the other side, 34,925 common hypo-methylated promoters (**Fig 3A**, blue square, **Suppl Tab 3B**) were strongly enriched in GO annotations referring to development (morphogenesis, axis specification, organ development and cell differentiation), germline and housekeeping functions (transcription, translation, metabolism, DNA repair, cell cycle, proliferation, etc). Some signaling, response to stimulus and transport terms also appeared enriched in this category. These data reveal that many genes supporting early development or housekeeping functions are by default in a permissive state for transcription. Even genes which are not necessarily required in the non-proliferative spermatozoa can nevertheless be primed for transcription in order to support embryo development. As an example, TSS region of *pola2*, a gene that encodes a subunit of the DNA polymerase, is equally hypomethylated in both tissues (**Fig 3B**). A vast majority of transcription factors instrumental for development *(otx, cbx* or *pax* families) also have constitutively hypo- or low-methylated promoters, even though they are not constitutively expressed. Their expression levels might be regulated by additional post-translational histone modifications as cells differentiate or face particular contexts.

Spermatozoa-specific hypo-methylated promoters belonging to 861 transcripts (**Fig 3A**, green shape, **Suppl Tab 3C**) were enriched in spermatozoa specific terms such as meiosis and germline, in addition to development, cellular metabolism and IGF signaling. Some of them are markers of the germline, such as *piwil1* involved in transposable elements silencing during gametogenesis, *dazl* which encodes a master translational regulator during spermatogenesis or the sex determining gene *dmrt1* (**Fig 3B**). Others, such as *spo11* or *mei4* encode specific meiotic recombination factors (**Fig 3B**). We also found the promoters of the pluripotency factors encoding *klf4* (**Fig 3B**) and *esrrb* to be specifically hypomethylated. Noticeably, a subset of developmental factors escapes constitutive hypomethylation and is specifically reprogrammed to be hypomethylated in the spermatozoa. This is the case of the entire *hox* clusters (**Fig 3B**), in which remarkably, altogether promoters: gene bodies and intergenic regions, spanning hundreds of kilobases are fully demethylated. The conserved *mir10b*, lying in the vicinity of *hox4* paralogs and co-expressed during development is therefore hypo-methylated as well. Interestingly *meis1a* (**Fig 3B**) or *meis3*, which are *hox* cofactors involved in axis patterning and organ morphogenesis are also specifically hypomethylated in the spermatozoa. This set of spermatozoa-specific genes is then silenced by methylation in the adult differentiated muscle, meaning that they might be sensitive to a dynamic regulation by promoter methylation.

Finally, 632 muscle-specific hypo-methylated promoters (**Fig 3A**, red shape, **Suppl Tab 3D**) were enriched in terms referring to terminal differentiation, muscle and vasculature function, cellular metabolism, response to stimulus and immune response. These dynamic promoters have undergone a demethylation during differentiation and are likely to be controlling muscle function. They mainly regulate terminal differentiation genes, such as *plecb* encoding a skeletal muscle plectin or signaling molecules such as *rarg* encoding a retinoic acid receptor (**Fig 3B**).

Altogether this work strongly supports the idea that sperm methylome is carrying an epigenetic program, which might be instrumental for both spermatogenesis and proper embryo development.

### Increasing rearing temperature alters sperm methylome

Given the importance of the paternal methylome in fish, we sought to question its robustness or sensitivity to environmental conditions. We chose to evaluate the potential impact of an increased rearing water temperature during gametogenesis on rainbow trout spermatozoa methylome. To this aim, we analyzed by WGBS the sperm of 6 fish bred at 16°C from May to November versus 6 control fish kept at 12°C (**Tab 1, Suppl Fig 1B**). Global average methylation rates were similar in both conditions (**Suppl Tab 1**) and the PCA analysis did not show any strong segregation between 12°C and 16°C spermatozoa samples (**Fig 4A and Suppl Fig 2**). Nevertheless, we identified 112,386 DMCs with a threshold adjusted p-value of 0.05 (**Fig 4B**). Nearly 50% of DMCs were hypomethylated (55,738) and 50% hypermethylated (56,648) at 16°C compared to 12°C. Interestingly, we found that 92% of these DMCs fall into differentially methylated regions (DMRs). This result indicates that an increase of rearing water temperature during gametogenesis alters spermatozoa methylome in a non-sporadic and non-random way. The affected cytosines instead concentrate into 5,359 DMRs of 400 bp median length resulting from a coordinated response to temperature increase. Ten % of these DMRs affect promoters, later on designed as prom-DMRs. Noteworthy, very few of these differentially methylated promoters had been previously identified as tissue specific, i.e. hypomethylated exclusively in either spermatozoa or muscle (11 and 7, respectively) (**Fig 5A**). On the contrary, they mostly fall in the common hypo-, common hyper- or common moderately-methylated promoter categories, already indicating that developmental genes epigenetically reprogrammed in the spermatozoa were mostly unaffected.

**Fig 4.**
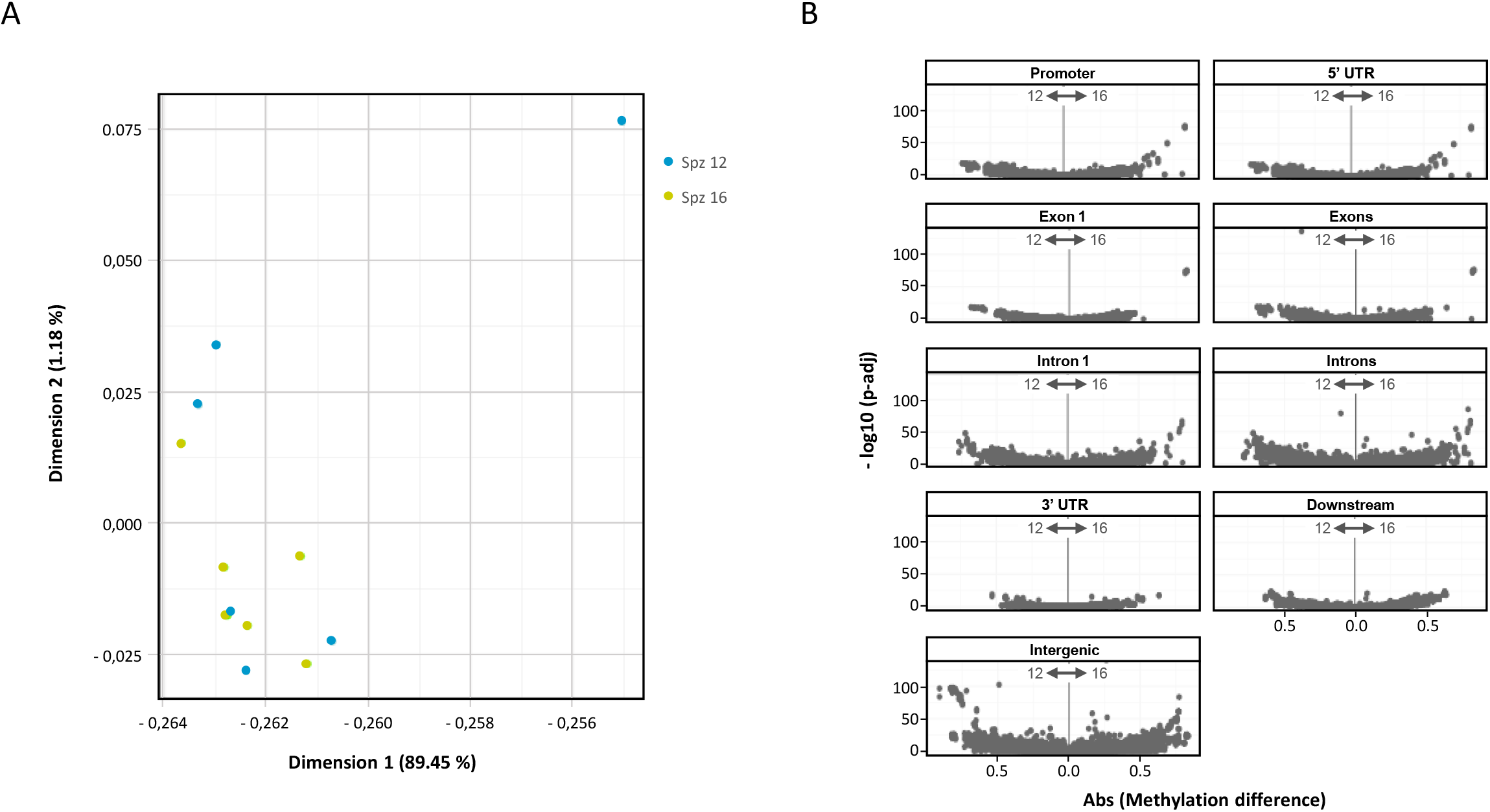
Differentially methylated cytosines between spermatozoa from males raised at 16°C versus 12°C. A. PCA plot of spermatozoa from males raised at 12°C and 16°C WGBS data sets. B. Volcano plots of DMCs between spermatozoa from males raised at 12°C versus 16°C in every genomic feature (× axis: absolute methylation difference, y: −log10 (adusted p-value)).

**Fig 5:**
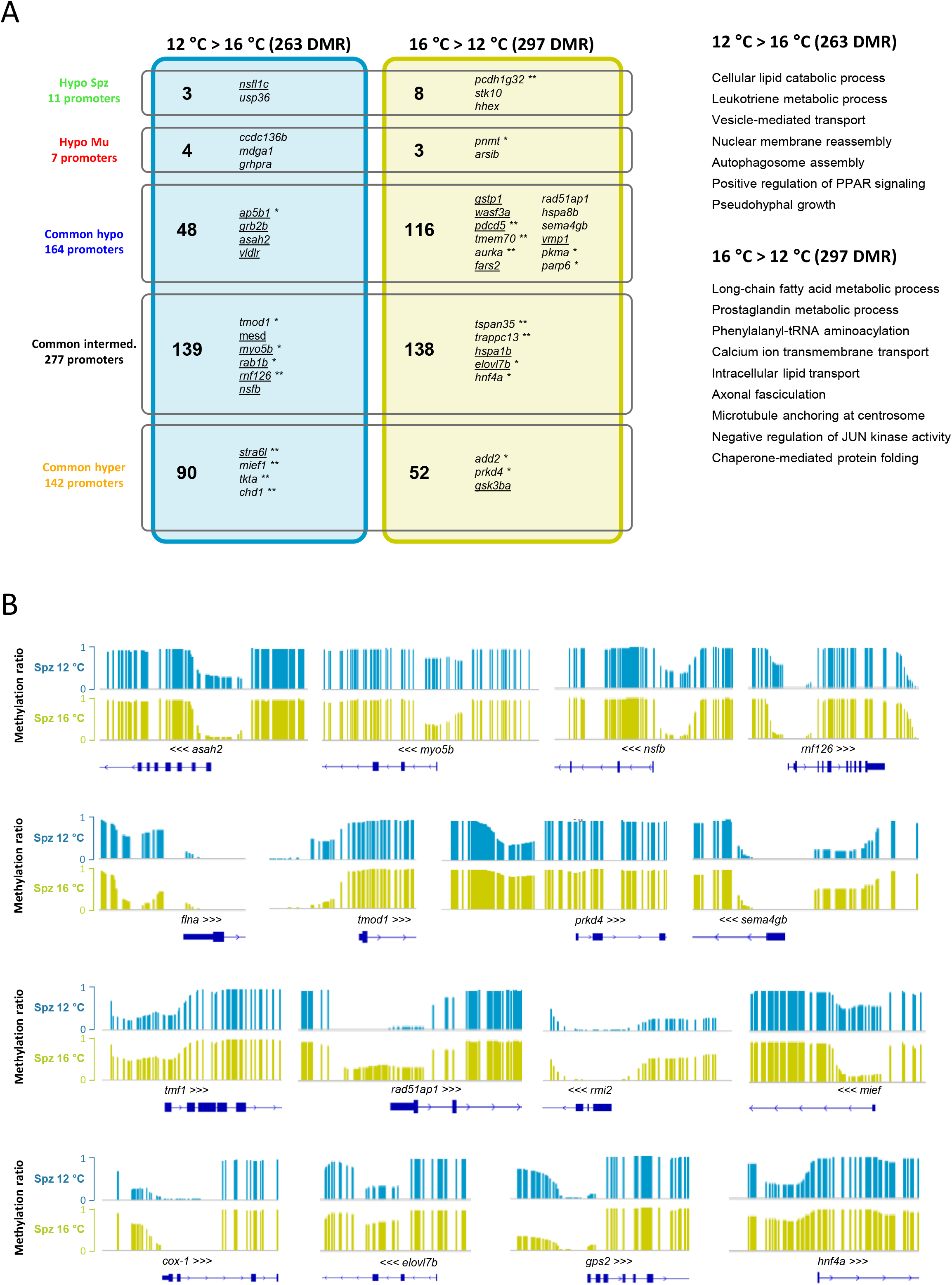
Differentially methylated gene promoters between spermatozoa of fish raised at 16°C *versus* 12°C. A. Intersection of up- and down-regulated DMRs with the different promoter categories defined in Fig 3 (sperm-specific hyper or hypo-methylated, equally methylated,…) and associated GO term enrichment analysis. Genes are given of examples of each categories, chosen either for their association to an enriched GO term or for their presence in the top200 list of prom-DMRs. underlined: associated to an enriched GO terms,*: top 10, **: top 50 prom-DMR. B. IGV snapshots of TSS regions as examples of hyper- and hypo-methylated promoters at 16°C versus 12°C.

In order to thoroughly study the biological functions of the differentially methylated promoters (prom-DMRs), we carried out both (i) a non-biased search for GO terms enrichment, (ii) a manual examination and annotation of the TOP 200 differentially methylated ones. Both approaches were mainly based on gene orthologies and mammalian literature. Examples of hypo- and hyper-methylated gene promoters and enriched GO categories are indicated in **Fig 5A** (**Suppl Tab 5A-B**). Interestingly, the manual annotation analysis largely confirmed the GO term enrichment test and indicated that the prom-DMRs impacted spermiogenesis and lipid metabolism genes, leaving most developmental genes unaffected.

#### Intracellular vesicle trafficking

We found prom-DMRs annotations covering virtually the whole spectrum of the molecular components regulating vesicular trafficking and/or autophagy: from plasma and organelle membrane regulation to motor proteins and signaling (**Fig 5A**, **Suppl Tab 5**). We found DMRs in the promoters of *asah2* (N-AcylSphingosine AmidoHydrolase) (**Fig 5B**), *ppm1la* and *cert1* (ceramide transporters), *sphk1* (sphingosine kinase) and *mdga1* (MAM Domain containing Glycosylphosphatidylinositol Anchor), which regulate membrane ceramid and sphingolipid metabolism, therefore affecting membrane properties, lipid rafts, protein sorting, signaling, plasma membrane curvature and endocytosis initiation (Pepperl et al., 2013; Tani et al., 2007; Young and Wang, 2018). The promoter of the membrane receptor *s1pr3* (protein-G-coupled receptor of Phospho-Sphingosine) is differentially methylated as well as the promoters of *vldlr* and *mesd* (very low density lipoprotein receptor and chaperone for low density lipoprotein receptors (LRP), respectively), likely affecting endocytosis initiation. We found several promoters of signaling molecules regulating vesicular traffic such as *grb2b, praf2* and *sh3gl1b* (adaptors), *rab1b* and *rnd2* (small GTPases), *dennd5a, rgl2, arhgef18a, quo, fgd4b* and *trappc13* (GEF, guanine nucleotide exchange factors for small GTPases), and *git2a*, *tbc1d2* and *rgs3b* (GAP, GTPases activating proteins). DMRs also affected promoters of vesicular traffic effectors such as the motor myosin *myo5b* (**Fig 5B**), the actin interacting WASP family member *wasf3a* and the kinesin *kif26ba*. Finally, we found prom-DMRs potentially affecting the regulation of membrane fusion, such as the promoters of *nsfl1c (p47* ortholog) and *nsfb (nsf* ortholog) (**Fig 5B**) encoding representative actors of the two major membrane fusion pathways (NSF and VCP/p47 pathways) (Brunger and DeLaBarre, 2003), the promoter of *rnf126* (E3 ubiquitin ligase, which is essential for proteasomal degradation in the VCP/p47 pathway) (**Fig 5B**), or the promoter of *myofl* involved in membrane fusion and repair. Markers of various types of intracellular vesicles where found differentially methylated, such as *ap5b1* and *ap5s1*, encoding two subunits of the late endosome AP-5 adaptor complex, *trappc13* encoding a component of the trafficking particle protein complex and *vmp1* (vacuole membrane protein), both involved in secretion and autophagy, therefore suggesting that several pathways orchestrating cellular trafficking might be elicited in response to increased rearing temperature. Therefore, sperm cells exposed to increased temperature exhibit a differential methylation at numerous promoters regulating vesicular trafficking potentially involving protein folding and degradation, cell signaling, autophagy or apoptosis.

#### Cytoskeleton remodeling

We found abundant prom-DMRs indicative of a reorganization of the cytoskeleton, mostly affecting actin network, cell-cell and cell-extra cellular matrix (ECM) adhesion (**Fig 5A**, **Suppl Tab 5**). As such, we found promoters of genes controlling actin dynamics: *flna* (filamin a) (**Fig 5B**), *tmod1* (tropomodulin) (**Fig 5B**), *add2* (adducin), *coro1b* (coronin) and *prkd4* (kinase) (**Fig 5B**), are all reported to interact with actin network and/or control its assembly, filament elongation or depolymerization. In addition, *sema4gb*, encoding a semaphorin was found among the prom-DMRs (**Fig 5B**). Semaphorins regulate intra-cellular actin filaments and microtubule network interactions with the ECM. Interestingly, in the same pathway, the promoters of the semaphorin receptors *plkdc2* (plexin) and *nrp1a* (neurpilin) and its co-factor *rftn2* (raftlin) were also differentially methylated. Other promoters of genes encoding components of cellular junctions or cell-ECM adhesion complexes were found among DMRs, such as the protocadherins *pcdh1g32* and *pcdhb*, and the cadherin interacting angiomotin-like *amotl2a* (adherens junctions), *nectin1a* (adherens junctions), the claudin encoding *cldnd* and *cldn23a* (tight junctions). Finally, other prom-DMRs were reported in the literature to be regulators of cell-cell and cell-ECM adhesion, such as *susd2*, *mpp3b*, *sh3bgrl*, *tbc1d2*, *mdga2a*, *cadm1b* and *rxylt1*. These results highlight a high occurrence of genes related to actin cytoskeleton remodeling, cell-cell and cell-ECM adhesion, among promoters affected by an increased temperature in spermatozoa. This suggests a potential response of differentiating germ cells at the level of both intra- and inter-cellular processes.

Therefore, numerous prom-DMRs annotations converge toward a functional adaptation of germ cells to a temperature stress with a coordinated cytoskeletal reorganization supporting an important modulation of intracellular vesicular trafficking and autophagy. This could be interpreted as a conserved and generic cytoprotective response to heat stress. Similar results were indeed reported by Madeira *et al*. who observed that larvae of sea bream *Sparus aurata* modulate, at the protein level, the intracellular transport, protein folding and degradation and the cytoskeleton dynamics pathways in response to temperature stress (Madeira et al., 2016). Indications of cytoskeleton remodeling in adaptation to increased temperature were also observed in a blue mussel gills (Fields et al., 2012) or in the myofibril tissue of the crab *P. cinctipes* (Garland et al., 2015). During their progression, male germ cells in rainbow trout thus modulate promoters classically involved in somatic heat stress adaptation and keep traces of these alterations in the terminally differentiated spermatozoa.

#### Spermiogenesis

Beyond the activation of possibly pan cell type adaptive mechanisms to increased rearing temperature, the cellular trafficking and cytoskeleton actors differentially methylated in our study interestingly pinpoint one major function precluding the spermatozoon stage: spermiogenesis. In the testis, differentiating sperm cells undergo drastic cell shape and cell-cell adhesion remodeling. These processes result from a tight control of the cytoskeleton dynamics and a recycling of intercellular junctions and ECM-interacting receptors at the plasma membrane, processes which are in turn dependent on endocytosis, vesicle trafficking, autophagy and protein recycling. Indeed, proper Golgi orientation, cytoskeleton, membrane fusion and vesicular trafficking are essential for the formation of the acrosome. Rainbow trout spermatozoa possess a transitory pseudo-acrosomal vesicle which is visible at the round spermatid stage and only leaves a cytoskeletal scar at later stages, reminiscent of mammalian acroplaxome (Billard, 1983). Endocytosis and vesicle trafficking are also crucial to the intraflagellar vesicle transport of molecules allowing the growth of the flagellum, and later on, for the elimination of the cytoplasm in late spermiogenesis and the individualization of sperm cells from the syncytium for spermiation. These spermiogenetic processes might recruit and coordinate common actors of vesicular trafficking expressed during spermiogenesis. It is therefore tempting to speculate that several gene promoters instrumental for spermiogenesis have been targeted by differential methylation. Interestingly, we found *tmf1* in the list of differentially methylated promoters (**Fig 5B**). Its mouse ortholog TMF/ARA160 encodes a protein associated to Golgi, emerging vesicles and microtubule network, which KO in male germ cells induces Golgi misorientation, lack of homing of pro-acrosomal vesicles, agenesis of the acrosome, inefficient cytoplasm removal and misshapen sperm head (Elkis et al., 2015; Lerer-Goldshtein et al., 2010). Myosin VI and Myosin Va, orthologs of *myo5b*, which we found differentially methylated in our data set, are essential for acrosome formation in the mouse and in the Chinese mitten crab (*E. sinensis*), respectively (Sun et al., 2010; Zakrzewski et al., 2020). The small GTPase *rnd2* mentioned earlier (“intracellular vesicle trafficking section”) was detected in Golgi and pro-acrosomal vesicles of rat spermatids (Naud et al., 2003).

In addition, several promoters of genes specifically implicated in other aspects spermatogenesis were found differentially methylated. As such, *rad51ap1* (**Fig 5B**), *rec8b* and *rmi2* (**Fig 5B**) encode essential actors of the meiotic homologous recombination (Bommi et al., 2019; Dray et al., 2011; Pires et al., 2017; Velkova et al., 2021), *aurka* encodes the aurora A kinase, involved in centrosome maturation and spindle formation during mitosis and also found to be essential for meiosis and spermiogenesis in mice (Lester et al., 2021), and *mief* (**Fig 5B**) encodes a mitochondrial fusion factor reminiscent of the mitochondrial remodeling to which rainbow trout spermatozoa are subjected during their differentiation (Zhao et al., 2011). Interestingly, while autophagy is involved in spermiogenesis, and in particular in acrosome formation (Wang et al., 2014), its increase in the human pathological model of cryptorchidism is associated to an impairment of spermatozoa maturation (Yefimova et al., 2019). Remarkably, in spite of the wide variation in spermatozoa morphologies and action modes among animals, cytoskeleton remodeling, vesicle trafficking and autophagy seem to be part of common pathways regulating normal, adaptive or pathological male germ cell maturation (White-Cooper and Bausek, 2010). Prom-DMRs which we detected in our experimental contrast are enriched in these GO categories. Altogether, these results argue in favor of an altered regulation of the late spermatogenic program (meiotic and post-meiotic) upon increased temperature. This could suggest that regions of the genome which were active during the exposure of the fish to the temperature stress (orchestrating late spermatogenesis) were more particularly prone to undergo a modulation of their methylation level.

#### Lipid metabolism

Interestingly, a third category of abundant prom-DMR annotations relates to the regulation of lipid metabolism (**Fig 5A**, **Suppl Tab 5**). We found several promoters of lipid catabolism genes such as *asah2* (ceramidase), *daglb* (lipase) or *abcd3a* (encoding a transporter involved in peroxisomal transport or catabolism of very long chain fatty acids). The regulation of long chain fatty acid metabolism was more specifically affected with the differential promoter methylation of *cox-1* (cyclooxygenase involved in arachidonic acid conversion) (**Fig 5B**), *ptges* (terminal enzyme of the cox-2-mediated prostaglandin E2 biosynthesis from arachidonic acid), *gstp1* (glutathione S-transferase), *gpx1* (glutathione peroxidase) and *elovl7b* (fatty acid elongase) (**Fig 5B**). In addition, the promoter of *gps2* (**Fig 5B**), which encodes a PPARɣ transcriptional co-activator, was found differentially methylated. Fatty acid composition in testis was shown to be of particular relevance for sperm maturation and function in human (Collodel et al., 2020), in ruminant (Van Tran et al., 2017), or rooster (Teymouri Zadeh et al., 2020). PPARɣ, known to be master regulator of adipocyte differentiation and lipid metabolism, was also shown to control fatty acid metabolism in human and therefore suspected to be instrumental for proper sperm formation (Olia Bagheri et al., 2021). Finally, the promoter of *hnf4a*, which controls fatty acid oxidation and metabolism in hepatocytes, was overlapping with a DMR in our analysis (**Fig 5B**). The appearance of fatty acid metabolism as a potentially altered pathway, controlled by PPARɣ and/or Hnf4a, could be interpreted as a metabolic adaptation of poikilotherm fish to temperature. The activity of several desaturases, elongases and fatty acid composition where indeed shown to be sensitive to temperature in fish and rainbow trout in particular (Tocher et al., 2004). It is remarkable however to find its scar in the methylation profile or spermatozoa.

## CONCLUSION

In the present study, we aimed at characterizing the rainbow trout sperm methylome and its sensitivity to a rearing temperature increase of 4°C during spermatogenesis. Single-base resolution methylomes revealed that an important pool of gene promoters is methylation-free in spermatozoa, and therefore in a permissive state for transcription, although sperm DNA is tightly packed and transcriptionally silent. Sperm cells thus carry an epigenetic code seemingly resulting from history and preparing their future role. Remarkably, sperm methylation-free promoters control housekeeping, early development and germline functions.

We found that an increased rearing temperature during spermatogenesis significantly impacted sperm methylome. Interestingly, DMRs affected promoters controlling spermiogenesis and lipid metabolism genes, leaving most developmental genes unaffected. This organized response to heat stress suggests a coordination by a signaling pathway. Our data highlighted PPARɣ and Hnf4 pathways. Alternatively, some genomic regions might be more prone to undergo methylome alterations because of a pre-established sensitivity, laying in the fact that they are active during spermiogenesis, which is both a period of deep chromatin remodeling and the time of experimental stress exposure. In an attempt to identify a potential driver of the differential methylation we observed, we looked for enriched transcription factor biding sites (TFBS) in our DMR using MEME suite. Interestingly, the most significantly enriched TFBS was the one of PRDM9 (**Suppl Fig 3**), which binds to recombination hotspots during meiosis. This opens the seducing possibility that meiotic recombination sites are at particular risk of epigenetic reprogramming upon environmental stress. Taken together, for the first time in rainbow trout, our results demonstrate that the methylation status of sperm-specific gene promoters controlling housekeeping and developmental function is very robust in the context of a 4°C temperature increase during spermatogenesis. Remarkably however, we found that the sperm methylome is altered and carry the traces of the life history of the individuals. We found epigenetic alterations of spermiogenesis and lipid metabolism controlling genes, which will be transmitted to the next generation. Future investigation should answer whether this altered sperm epigenome could be a molecular basis of acclimation to a heat stress for next generations and whether it could impact positively or negatively offspring performances under various conditions.

## MATERIAL AND METHODS

### Ethics statements

The experiment was conducted following the Guidelines of the National Legislation on Animal Care of the French Ministry of Research (Decree No 2013–118, February 2013) and in accordance with EU legal frameworks related to the protection of animals used for scientific purposes (i.e., Directive 2010/63/EU). The scientists in charge of the experiments received training and personal authorization. The experiment was conducted at the INRAE Physiology and Genomic Laboratory (LPGP) experimental facilities (permit number C35-238-6, Rennes, France), and approved by the ethics committee for the animal experimentation of Rennes under the authorization number T-2019-05-AL.

### Fish rearing and sampling

*Oncorhynchus mykiss* rainbow trouts were reared in our experimental farm and transferred to our indoor fish facility in April of their second year. Two groups of 20 males were kept in a recycling system, under artificial spring/summer/autumn photoperiod, and at the temperatures of 12°C and 16°C (after 2 weeks acclimation and gradual temperature increase). Growth of the fish in both groups were recorded and their commercial diet rations were adapted so that both groups follow similar growth curves. Fish sperms were harvested by stripping and muscle samples were taken after euthanasia (tricaine 200 mg/mL) when they reached the reproduction period.

### Whole genome bisulfite sequencing

Genomic DNA was prepared after an overnight lysis in TNES/Urea buffer (10 mM Tris, 0.125 M NaCl, 10 mM EDTA, 17 mM SDS, 4M urea, pH 8) complemented with 80ug/mL proteinase K at 42°C. After phenol-chloroform extraction, the samples were treated with RNase A (4mg/mL) and gDNAs were precipitated by the addition of isopropanol and sodium acetate. Whole genome bisulfite sequencing libraries were prepared (according to Accel-NGS Methyl-Seq DNA library Kit for Illumina protocol, Swift Biosciences) and sequenced in 150 bp paired-end reads by Novogene company.

### Bio-informatic analysis

Quality and adapter trimming of the reads was carried out using Trim Galore (with an additional 10 bp clipping for read2 5’-end and read1 3’-end) (Krueger and Galore, n.d.). Trimmed reads mapping on Omyk_1.0 genome and methylation calls were performed with Bismark (F. Krueger and Andrews, 2011). BigWig files were obtained with bedGraphToBigWig (Kent et al., 2010) and methylation values were visualized with Integrative Genome Viewer (Robinson et al., 2011). These steps were included in a workflow which agrees to FAIR principles and is accessible online (https://forgemia.inra.fr/lpgp/methylome). The methylome (v1.0) workflow was built in Nexflow dsl2 language using a singularity container and optimized in terms of computing resources (cpu, memory) for its use on an informatic farm with a slurm scheduler.

Differentially methylated cytosines (DMCs) were identified by the R package DSS (Feng et al., 2014; Wu et al., 2015), using the optional methylation level smoothing on 500 bp and a Benjamini-Hochberg adjusted p-value threshold of 0,05. Differentially methylated regions (DMRs) were defined using DSS package with seed regions containing at least 5 CpGs and expansion window criterias agreeing with a minimum of 75 % of DMCs (with raw pval < 0,01).

CpGs were annotated with GenomeFeatures (Brionne, n.d.), RepeatMasker (Smit, 2015) and ensembl v105.1 annotation. Biological interpretations were carried out using ViSEAGO R package (Brionne et al., 2019) and the Gene Ontology (GO) public database. Associated gene terms were implemented from the extended Ensembl Compara custom annotation based on the release 103. Enrichment tests were performed using exact Fisher’s test. Enriched GO terms (p-val < 0,01) were grouped into functional clusters by Wang’s semantic similarity respecting GO graph topology and Ward’s criterion.

## ACKNOWLEDGEMENTS

The study was funded by the department of animal physiology (PHASE) of INRAE (ACI 2019). We thank Julien Bobe for its contribution to the experimental design and his comments on the manuscript. We thank the staff of our experimental farm (PEIMA, INRAE) and fish facility (ISC LPGP) and particularly Cécile Duret for dedicated handling of our experimental fish.

## AUTHORS CONTRIBUTIONS STATEMENT

AL designed the project and experiments, supervised the study and wrote the manuscript. CL and JJL contributed to the design of the project. ASG and CL performed the wet lab work. AB performed the bio-informatic analysis. All the authors read and approved the manuscript.

---

**Suppl Fig 1.**
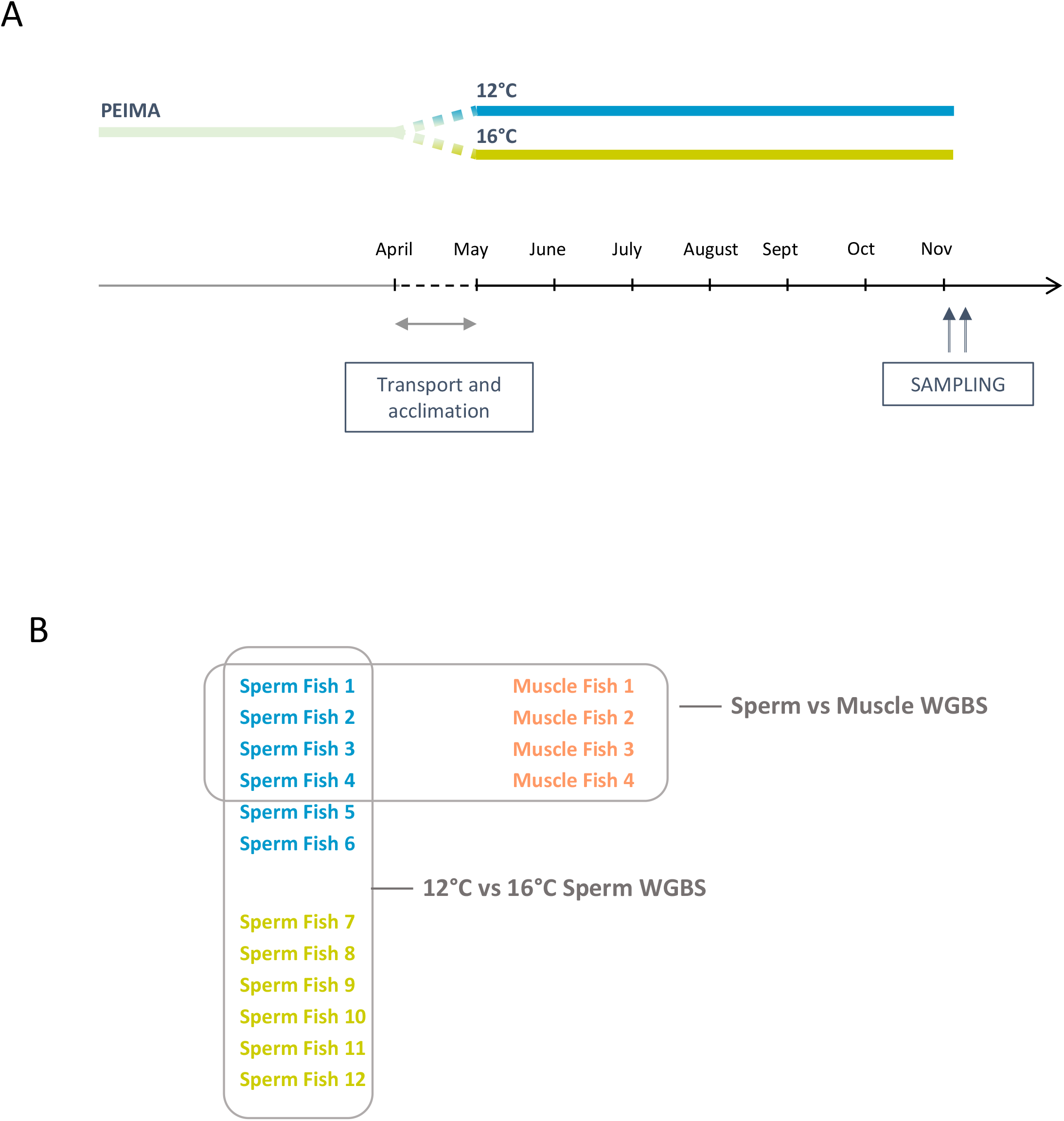
Experimental design. A. *Oncorhynchus mykiss* rainbow trouts were reared in our experimental farm and transferred to our indoor fish facility in April of their second year. Two groups of 20 males were kept in a recycling system, under artificial spring/summer/autumn photoperiod, and at the temperatures of 12°C and 16°C (after 2 weeks acclimation and gradual temperature increase). Fish sperm and muscle were sampled during the reproduction period. B. Four fish were sequenced for the sperm versus muscle paired analysis while two groups of 6 fish were sequenced for the study of the temperature contrast in spermatozoa.

**Suppl Fig 2.**
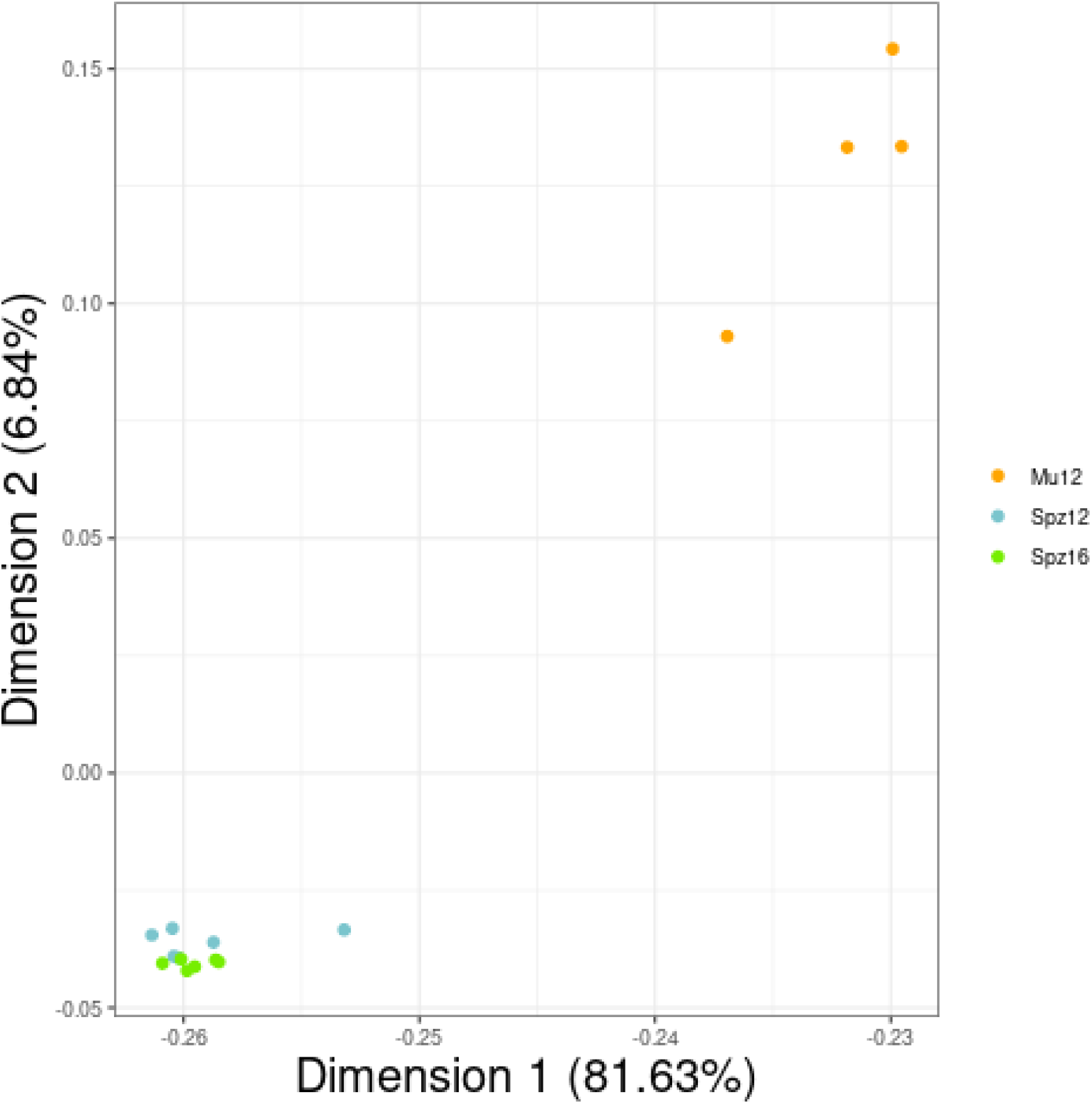
PCA plot of muscle (Mu12), spermatozoa from males raised at 12°C (Spz12) and spermatozoa from males raised at 16°C (Spz16) WGBS data sets.

**Suppl Fig 3.**
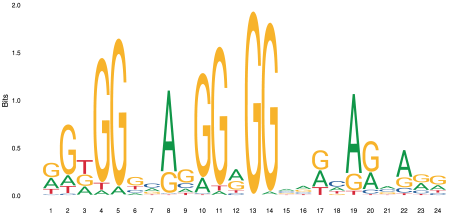
Putative recombination hotspots enrichment in DMRs between spermatozoa from males raised at 16°C versus 12°C. Vertebrate PRDM9 consensus DNA binding site identified by MEME in DMR sequences between spermatozoa from males raised at 16°C versus 12°C, and associated p-vamue. MEME suite is accessible online (https://meme-suite.org/meme/), SEA algorithm was used.

## Notes

### Competing Interest Statement

The authors have declared no competing interest.

