## Supplemental Figure 1 for "EPIGENETIC LANDSCAPE OF HEAT STRESS INTERGENERATIONAL INHERITANCE IN A TELEOST FISH"

### Supplementary Figure 1

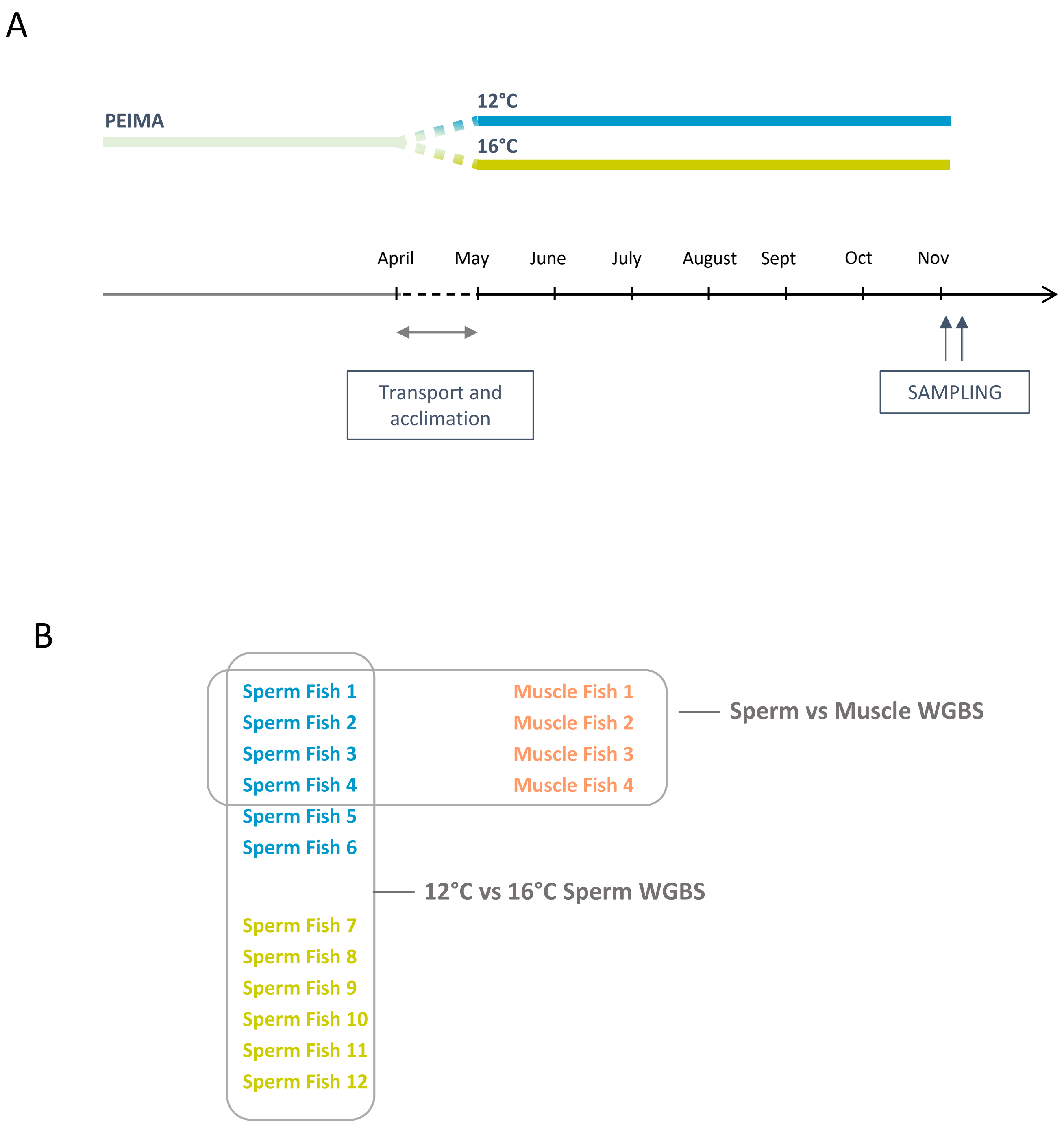


**Suppl Fig 1. Experimental design**

A. *Oncorhynchus mykiss* rainbow trouts were reared in our experimental farm and transferred to our indoor fish facility in April of their second year. Two groups of 20 males were kept in a recycling system, under artificial spring/summer/autumn photoperiod, and at the temperatures of 12°C and 16°C (after 2 weeks acclimation and gradual temperature increase). Fish sperm and muscle were sampled during the reproduction period. B. Four fish were sequenced for the sperm versus muscle paired analysis while two groups of 6 fish were sequenced for the study of the temperature contrast in spermatozoa.
