## Supplemental Figure 2 for "EPIGENETIC LANDSCAPE OF HEAT STRESS INTERGENERATIONAL INHERITANCE IN A TELEOST FISH"

Supplementary Figure 2

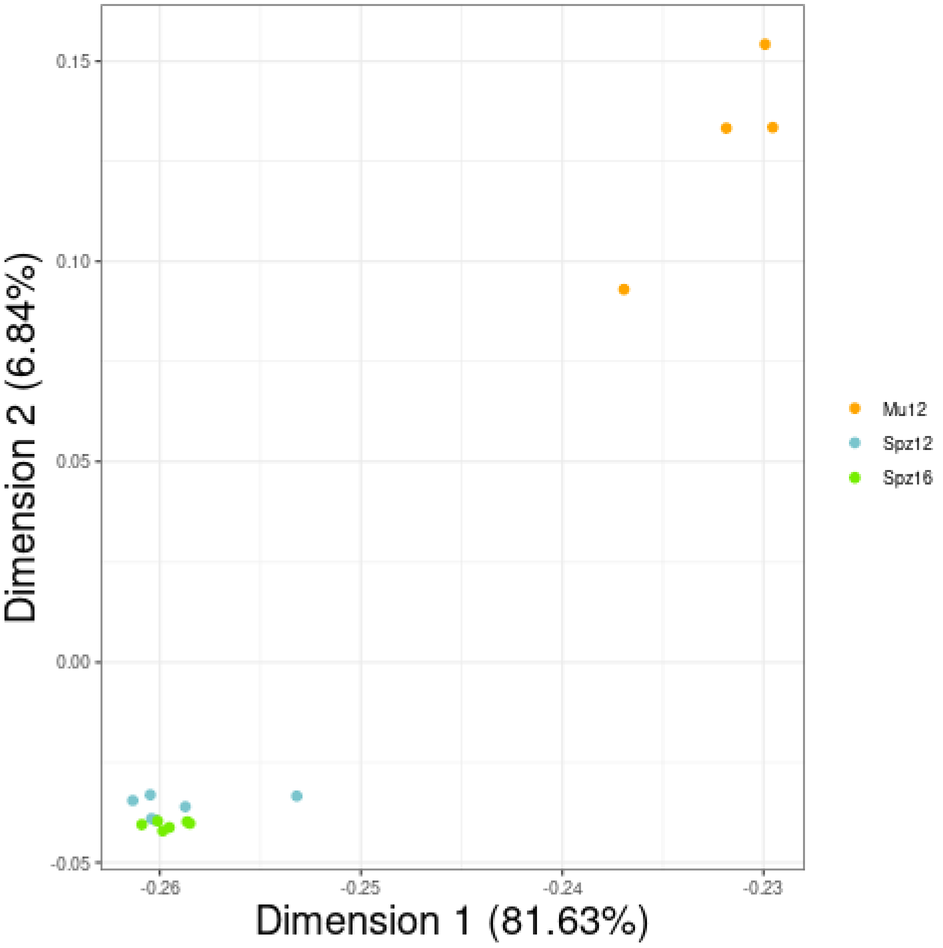


**Suppl Fig 2. PCA plot of muscle (Mu12), spermatozoa from males raised at 12°C (Spz12) and spermatozoa from males raised at 16°C (Spz16) WGBS data sets**
