## Supplemental Figure 3 for "EPIGENETIC LANDSCAPE OF HEAT STRESS INTERGENERATIONAL INHERITANCE IN A TELEOST FISH"

### Suppl Fig 3

PRDM9 p-val 9.28e-35

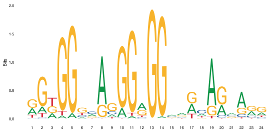

**Suppl Fig 3. Putative recombination hotspots enrichment in DMRs between spermatozoa from males raised at 16°C versus 12°C.**

Vertebrate PRDM9 consensus DNA binding site identified by MEME in DMR sequences between spermatozoa from males raised at 16°C versus 12°C, and associated p-vamue.

MEME suite is accessible online (https://meme-suite.org/meme/), SEA algorithm was used.
